# Retrospective Association Analysis of Longitudinal Binary Traits Identifies Important Loci and Pathways in Cocaine Use

**DOI:** 10.1101/628180

**Authors:** Weimiao Wu, Zhong Wang, Ke Xu, Xinyu Zhang, Amei Amei, Joel Gelernter, Hongyu Zhao, Amy C. Justice, Zuoheng Wang

## Abstract

Longitudinal phenotypes have been increasingly available in genome-wide association studies (GWAS) and electronic health record-based studies for identification of genetic variants that influence complex traits over time. For longitudinal binary data, there remain significant challenges in gene mapping, including misspecification of the model for the phenotype distribution due to ascertainment. Here, we propose L-BRAT, a retrospective, generalized estimating equations-based method for genetic association analysis of longitudinal binary outcomes. We also develop RGMMAT, a retrospective, generalized linear mixed model-based association test. Both tests are retrospective score approaches in which genotypes are treated as random conditional on phenotype and covariates. They allow both static and time-varying covariates to be included in the analysis. Through simulations, we illustrated that retrospective association tests are robust to ascertainment and other types of phenotype model misspecification, and gain power over previous association methods. We applied L-BRAT and RGMMAT to a genome-wide association analysis of repeated measures of cocaine use in a longitudinal cohort. Pathway analysis implicated association with opioid signaling and axonal guidance signaling pathways. Lastly, we replicated important pathways in an independent cocaine dependence case-control GWAS. Our results illustrate that L-BRAT is able to detect important loci and pathways in a genome scan and to provide insights into genetic architecture of cocaine use.

## 1. Introduction

Genome-wide association studies (GWAS) have successfully discovered many disease susceptibility loci and provided insights into the genetic architecture of numerous human complex diseases and traits. In some genetic epidemiological studies, longitudinally collected phenotype data are available. This is the case for many electronic health record (EHR)-based studies. As many of these studies continue to follow enrolled subjects (e.g. the UK Biobank (UKB) and the Million Veteran Program (MVP)), longitudinal phenotypes will be increasingly available with the passage of time, providing new data resources that require appropriate analytical tools for optimal analysis. Standard association tests that consider one time point or collapse repeated measurements into a single value such as an average do not capture the trajectory of phenotypic traits over time and may result in a loss of statistical power to detect genetic associations. In addition, the effects of time-varying covariates cannot be easily incorporated in such analyses. Recently, methodological developments for GWAS have proliferated to make full use of the available longitudinal data. For population cohorts, methods that account for dependence among observations from an individual include mixed effects models (Furlotte et al. 2012; Sikorska et al. 2013), generalized estimating equations (GEE) (Sitlani et al. 2015), growth mixture models (Das et al. 2011; Londono et al. 2013), and empirical Bayes models (Meirelles et al. 2013). Most of these approaches are prospective analyses and have been successfully applied to quantitative phenotypes.

As many diseases are rare, efficient designs, such as the case-control design, are commonly applied in epidemiological studies to recruit study subjects. Despite the enhanced efficiency in the study sample, non-random ascertainment can be a major source of model misspecification that may lead to inflated type I error and/or power loss in association analysis. The linear mixed model and the logistic mixed model do not perform well when the case-control ratio is unbalanced in large-scale genetic association studies (Zhou et al. 2018). Prospective analysis in which a population-based model is used ignores ascertainment bias and can result in compromised statistical inference. Furthermore, in the ascertained sample, the prospective approach conditional on the genotype and covariates may lose information when the joint distribution of the genotype and covariates carries additional information on whether the phenotype is associated with the genotype (Jiang et al. 2015). In this regard, several retrospective association methods have been proposed for analyzing ascertained population-based case-control studies (Hayeck et al. 2015; Jiang et al. 2016), family-based studies of continuous traits (Jakobsdottir and McPeek 2013), family-based case-control studies (Zhong et al. 2016; Hayeck et al. 2017), and family-based longitudinal quantitative traits (Wu and McPeek 2018). Compared to prospective tests, retrospective tests conditional on the phenotype and covariates are more robust to misspecification of the trait model (Jiang et al. 2015).

To generalize case-control sampling, outcome-dependent sampling designs have become popular for binary data in longitudinal cohort studies (Schildcrout and Heagerty 2008; Schildcrout et al. 2018a,b). However, association tests for longitudinally measured binary data are less well developed in GWAS. Here, we propose L-BRAT, a retrospective, GEE-based method for genetic association analysis of longitudinal binary outcomes. It requires specification of the mean of the outcome distribution and a working correlation matrix for repeated measurements. L-BRAT is a retrospective score approach in which genotypes are treated as random conditional on the phenotype and covariates. Thus, it is robust to ascertainment and trait model misspecification. It allows both static and time-varying covariates to be included in the analysis. We note that GMMAT, a recently proposed prospective test using the logistic mixed model to control for population structure and cryptic relatedness in case-control studies (Chen et al. 2016), can be adapted for repeated binary data. For comparison, we also develop RGMMAT, a retrospective, generalized linear mixed model (GLMM)-based association test for longitudinal binary traits.

We performed simulation studies to evaluate the type I error and power of L-BRAT and RGMMAT, and compared them to the existing prospective methods. The results demonstrate that the retrospective association tests have better control of type I error when the phenotype model is misspecified, and are robust to various ascertainment schemes. Moreover, they are more powerful than the prospective tests. Finally, we applied L-BRAT and RGMMAT to a genome-wide association analysis of repeated measurements of cocaine use in a longitudinal cohort, the Veterans Aging Cohort Study (VACS), and replicated the results using data from an independent cocaine dependence case-control GWAS.

## 2. Methods

Suppose a binary trait is measured over time on a study population of *n* individuals. We have their genome-wide measures of genetic variation. A set of covariates, static or dynamic, are also available. Let *n_i_* be the number of repeated measures on individual *i* and 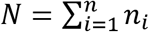 be the total number of observations. For individual *i*, let ***X**_ij_* and *Y_ij_* be the *p*-dimensional covariate vector, assumed to include an intercept, and the binary response at time *t_ij_*, respectively. In this setting, individuals are permitted to have measurements at different time points and different number of observations. We let ***Y*** denote the outcome vector of length *N*, and let ***X*** denote the *N* × *p* covariate matrix. Here, we focus on the problem of testing for association between a genetic variant and the longitudinal binary outcomes. Let ***G*** denote the vector of genotypes for the *n* individuals at the variant to be tested, where *G_i_* = 0, 1, or 2 is the number of minor alleles of individual *i* at the variant.

### 2.1. GEE-based Model

We consider a GEE approach in which the mean of the outcome distribution, given the genotype and covariates, is specified as

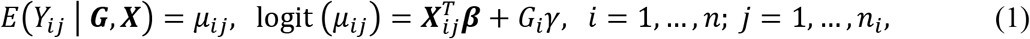

where ***β*** is a *p*-dimensional vector of covariate effects and *γ* is a scalar parameter of interest representing the effect of the tested variant. Writing in a matrix form, we have the mean model

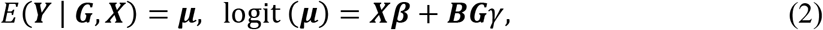

where ***B*** is an *N* × *n* matrix representing the measurement clustering structure, and its (*l, i*)th entry *B_li_* is an indicator of the *l*th entry of ***Y*** being a measurement on individual *i*. Here, the vector ***BG*** is the vertically expanded genotype vector that maps the genotype data ***G*** from the individual level to the measurement level. The covariance structure of ***Y*** is given by

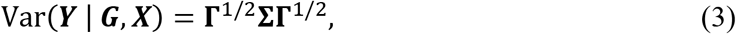

where **Γ** = diag{*μ*_1,1_(1 − *μ*_1,1_),…, *μ*_1,*n*_1__(1 − *μ*_1,*n*_1__),…, *μ*_*n*,1_(1 − *μ*_*n*,1_),…, *μ_n,n_n__*(1 − *μ_n, n_n__*)} is an *N*-dimensional diagonal matrix and **Σ** is an *N* × *N* correlation matrix. The covariance specification in Eq. (3) ensures that the variance of the dichotomous response *Y_ij_* depends on its mean in a way that is consistent with the Bernoulli distribution. To apply the GEE method, a working correlation structure such as independent, exchangeable, and first-order autoregressive (AR(1)) must be specified. For a given within-cluster correlation matrix **Σ**(*τ*), which may depend on some parameter *τ*, the estimating equations for the unknown parameters (***β**, γ*) are written as

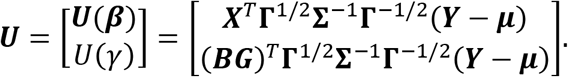

To detect association between the genetic variant and the phenotype, we consider a score approach to test *H*_0_:*γ* = 0 against *H*_1_: *γ* ≠ 0. The null estimate of ***β***, denoted by 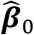, is the solution to a system of estimating equations ***U***(***β***) = 0 under the constraint *γ* = 0, which can be computed iteratively between a Fisher scoring algorithm for ***β*** and the method of moments for *τ* until convergence. Then, the score function for *γ* is

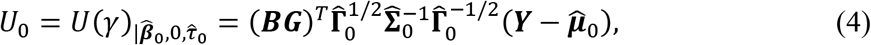

where 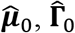 and 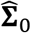 are ***μ*, Γ** and **Σ** evaluated at 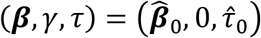.

In the GEE approach, the prospective score statistic for testing *H*_0_:*γ* = 0 takes the form

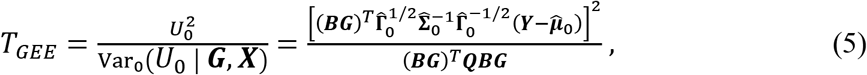

where the null variance of *U*_0_ is evaluated using a robust sandwich variance estimator, conditional on the genotype and covariates. Here ***Q*** = ***V*** − ***VX***(***X**^T^**VX***)^−1^***X***^*T*^***V***, where 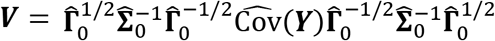 and the sample covariance of 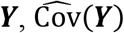, is estimated by 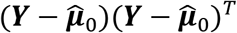. Under the null hypothesis, the *T_GEE_* test statistic has an asymptotic 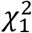 distribution.

In what follows, we introduce a new GEE-based association testing method, L-BRAT (Longitudinal Binary-trait Retrospective Association Test). The L-BRAT test statistic is also based on the score function *U*_0_ in Eq. (4). In contrast to the prospective GEE score test, L-BRAT takes a retrospective approach in which the variance of *U*_0_ is assessed using a retrospective model of the genotype given the phenotype and covariates. An advantage of the retrospective approach is that the analysis is less dependent on the correct specification of the phenotype model. We assume that under the null hypothesis of no association between the genetic variant and the phenotype, the quasi-likelihood model of ***G*** conditional on ***Y*** and ***X*** is

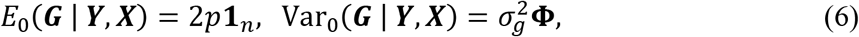

where *p* is the minor allele frequency (MAF) of the tested variant, **1**_*n*_ is a vector of length *n* with every element equal to 1, 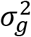 is an unknown variance parameter, and **Φ** is an *n* × *n* genetic relationship matrix (GRM) representing the overall genetic similarity between individuals due to population structure. Because ***B*1**_*n*_ = **1**_*N*_, which is the first column of ***X*** that encodes an intercept, and 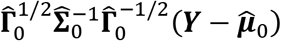, the *N*-dimensional vector of transformed null phenotypic residuals, is orthogonal to the column space of ***X***, then the null mean model of ***G*** in Eq. (6) ensures that

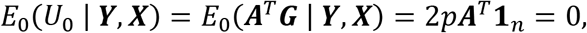

where 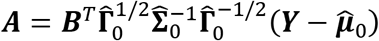 is the individual-level transformed phenotypic residual vector of length *n*.

In model (6), the GRM **Φ** can be obtained using genome-wide data, given by

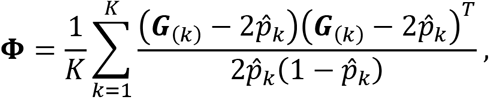

where *K* is the total number of genotyped variants, ***G***_(*k*)_ is the genotype vector at the *k*th variant, and 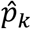 is the estimated MAF, for example, 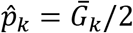, the sample MAF at the *k*th variant. For the variant of interest, let 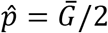 be its sample MAF. Under Hardy-Weinberg equilibrium, the variance of the genotype is estimated by 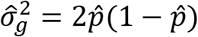. Or we can use a more robust variance estimator (Jakobsdottir and McPeek 2013) given by

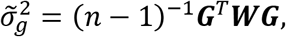

where 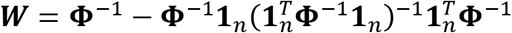. Finally, the L-BRAT test statistic can be defined as

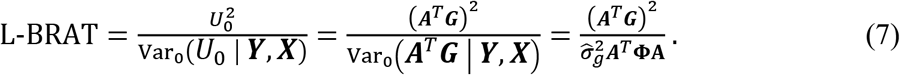

Under regularity conditions, L-BRAT asymptotically follows a 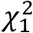 distribution under the null hypothesis.

### 2.2. GLMM-based Model

The Generalized linear Mixed Model Association Test (GMMAT) was originally designed to use random effects in logistic mixed models to account for population structure and cryptic relatedness in case-control studies (Chen et al. 2016). To extend the GMMAT method for case-control analysis to repeated binary data, we consider the following logistic mixed model:

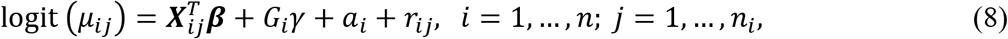

where *μ_ij_* = *P*(*Y_ij_* = 1 | *G_i_, **X**_ij_, a_i_, r_ij_*) is the probability of a binary response at time *t_ij_* for individual *i*, conditional on his/her genotype, covariates, and random effects *a_i_* and *r_ij_*, ***β*** and *γ* are the same as defined in model (1), *a_i_* is the individual random effect, and *r_ij_* is the individual-specific time-dependent random effect. The two random effects were used to capture the correlation among repeated measures in gene-based association test for longitudinal traits (Wang et al. 2017). Here, *a_i_*’s are assumed to be independent and 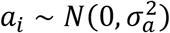. The vector of time-dependent random effects ***r**_i_* = (*r*_*i*1_,…, *r_i,n_i__*) has a multivariate normal distribution, 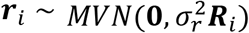, where an AR(1) structure is assumed for the correlation matrix ***R**_i_*, in which *τ* is the unknown parameter. The binary responses *Y_ij_* are assumed to be independent given the random effects *a_i_* and *r_ij_*.

To construct a score test for the null hypothesis *H*_0_: *γ* = 0 vs. the alternative *H*_1_: *γ* ≠ 0, we use the penalized quasi-likelihood method (Breslow and Clayton 1993) to fit the null logistic mixed model and obtain the null estimates of ***β***, 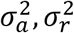 and *τ*, denoted by 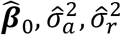 and 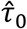 (Chen et al. 2016). Similarly, the best linear unbiased predictor (BLUP) of random effects, 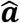 and 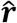, can be obtained. Then, the resulting score function for *γ* is

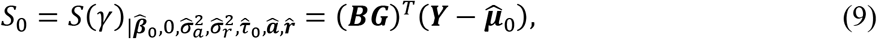

where 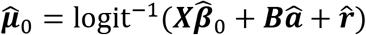 is a vector of fitted values under *H*_0_.

In GMMAT, the null variance of the score *S*_0_ is evaluated prospectively (Chen et al. 2016), i.e., Var_0_(*S*_0_ | ***G,X***) = (***BG***)^*T*^***PBG***, where ***P*** = **Ψ**^−1^ − **Ψ**^−1^***X***(***X***^*T*^**Ψ**^−1^***X***)^−1^***X***^*T*^**Ψ**^−1^, and 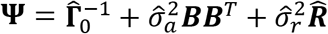. Here 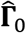 and 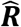 are **Γ** and ***R*** evaluated at 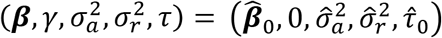, where **Γ** is the same as defined in Eq. (3) and ***R*** = diag{***R***_1_,…, ***R***_*n*_} is a block diagonal matrix. The GMMAT test statistic can be written as

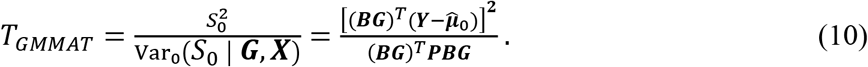

Like L-BRAT, we can construct a retrospective test to assess the significance of the GLMM score function of Eq. (9), which we call RGMMAT, based on the quasi-likelihood model of ***G*** in Eq. (6). Thus, we define the RGMMAT statistic by

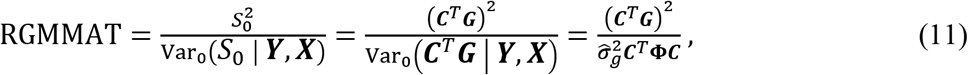

where 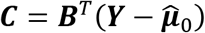 is the *n*-dimensional vector of phenotypic residuals at the individual level by summing over all time points for an individual, and the phenotypic residuals are obtained by fitting the null logistic mixed model. Both the GMMAT and RGMMAT test statistics are assumed to have 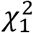 asymptotic null distributions.

## 3. Simulation Studies

We performed simulation studies to evaluate the type I error and power of the two retrospective tests we propose, and compared them to the prospective GEE and GMMAT methods. We also assessed sensitivity of L-BRAT and RGMMAT in the presence of model misspecification and ascertainment. In the simulations, we considered two different trait models and three different ascertainment schemes. Because both the L-BRAT and GEE methods require specification of a working correlation matrix, we implemented three working correlation structures: (1) independent, (2) AR(1), and (3) a mixture of exchangeable and AR(1).

To generate genotypes, we first simulated 10,000 chromosomes over a 1 Mb region using a coalescent model that mimics the linkage disequilibrium (LD) and recombination rates of the European population (Schaffner et al. 2005). We then randomly selected 1,000 non-causal single nucleotide polymorphisms (SNPs) with MAF > 0.05. In addition, we generated two causal SNPs that were assumed to influence the trait value with epistasis. In the type I error simulations, we tested association at the 1,000 non-causal SNPs. In each simulation setting, we generated 1,000 sets of phenotypes at five time points. Putting together, 10^6^ replicates were used for the type I error evaluation. In the power simulations, we tested the first of the two causal SNPs and empirical power was assessed using 1,000 simulation replicates. In all tests considered, the genotypes at the untested causal SNP(s) were assumed to be unobserved.

### 3.1. Trait Models

We simulated two types of binary trait models at five time points, in which the two unlinked causal SNPs were assumed to act on the phenotype epistatically. The first type is a logistic mixed model, given by

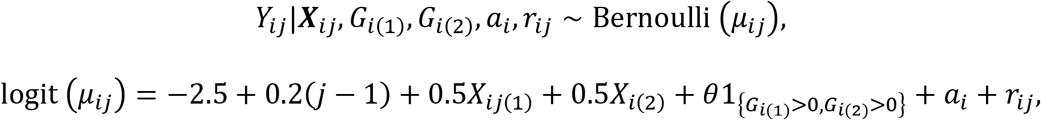

where *X*_*ij*(1)_ is a continuous, time-varying covariate generated independently from a standard normal distribution, *X*_*i*(2)_ is a binary, time-invariant covariate taking values 0 or 1 with a probability of 0.5, *G*_*i*(1)_ and *G*_*i*(2)_ are the genotypes of individual *i* at the two causal SNPs, *θ* is a scalar parameter encoding the effect of the causal SNPs, 1_{*G*_*i*(1)_>0,*G*_*i*(2)_>0}_ is an indicator function that takes value 1 when individual *i* has at least one copy of the minor allele at both the causal SNPs, *a_i_* and *r_ij_* are the individual-level time-independent and time-dependent random effects, respectively. Here we assume 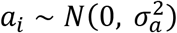 and 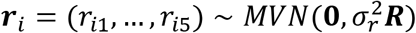, where ***R*** is a 5 × 5 correlation matrix specified by the AR(1) structure with a correlation coefficient *τ*. The two causal SNPs are assumed to be unlinked with MAFs 0.1 and 0.5, respectively. The variance components are set to 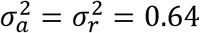 and *τ* = 0.7.

The second type of trait model we considered is a liability threshold model in which an underlying continuous liability determines the outcome value based on a threshold. Specifically, the phenotype *Y_ij_* is given by

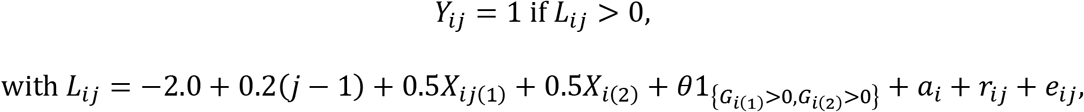

where *L_ij_* is the underlying liability for individual *i* at time *t_ij_*, and 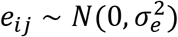 represents independent noise, with 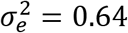. All other parameters are the same as those in the logistic mixed model.

In both models, we included a time effect and assumed that the mean of the outcome increases with time. The effect of the causal SNPs was set to *θ* = 0.34 in the type I error simulations. For the power evaluation, we considered a range of values for *θ*, where we set *θ* = 0.3, 0.32, 0.34, 0.36, and 0.38. At the given parameter values, the prevalence of the event of interest ranges from 12.8% to 27.7% over time. The proportion of the phenotypic variance explained by the two causal SNPs ranges from 0.69% to 1.10% in the logistic mixed model, and from 0.49% to 0.78% in the liability threshold model.

### 3.2. Sampling Designs

We considered three different sampling designs. In the “random” sampling scheme, the sample contains 2,000 individuals that were randomly selected from the population regardless of their phenotypes. Thus, ascertainment is population based. In the “baseline” sampling scheme, we sampled 1,000 case subjects and 1,000 control subjects according to their outcome value at baseline only. In the “sum” sampling scheme, individuals were stratified into three strata (1, 2, and 3) based on a total count that sums each subject’s response over time, where samples in stratum 1 never experienced the event of interest, i.e., 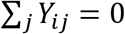, samples in stratum 2 sometimes experienced the event, i.e., 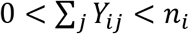, and samples in stratum 3 always experienced the event, i.e., 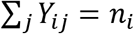. Following the outcome-dependent sampling design for longitudinal binary data (Schildcrout et al. 2018b), we selected 100, 1,800, and 100 individuals from the three strata respectively to oversample subjects who have response variation over the course of the study.

### 3.3. Simulation Results

To assess type I error, we tested for association at unlinked and unassociated SNPs. Table 1 gives the empirical type I error of the L-BRAT, RGMMAT, GEE, and GMMAT tests, based on 10^6^ replicates, at the nominal type I error level *α*, for *α* = 0.05, 0.01, 0.001, and 0.0001. In all simulations, the type I error of the two retrospective tests, L-BRAT and RGMMAT, exhibited no inflation at any of the nominal levels considered. In contrast, the prospective GEE tests, regardless of the choice of working correlation, had inflated type I error at most of the nominal levels in all settings. This is likely due to the fact that the asymptotic distribution of robust sandwich variance estimators used in GEE are not well calibrated. The inflated type I error was also reported in longitudinal GWAS with quantitative traits using GEE (Sitlani et al. 2015). In GMMAT, the type I error was much lower than the nominal level when *α* = 0.05, 0.01, 0.001, and 0.0001. These results demonstrate that the two retrospective tests, L-BRAT and RGMMAT, are robust to trait model misspecification and ascertainment, whereas GEE has type I error inflation and GMMAT is overly conservative. Overall, the choice of the working correlation matrix does not have much impact on the type I error of the L-BRAT method.

**Table 1.** Empirical type I error of L-BRAT, RGMMAT, GEE, and GMMAT, based on 10^6^ replicates. Rates that are significantly larger than the nominal levels are in bold. Texts in the brackets following test statistics denote the working correlation structure. Specifically, L-BRAT(ind) and GEE(ind) denote the L-BRAT and GEE tests with an independent working correlation; L-BRAT(AR1) and GEE(AR1) denote the L-BRAT and GEE tests with an AR(1) working correlation; L-BRAT(mix) and GEE(mix) denote the L-BRAT and GEE tests with a mixture of exchangeable and AR(1) working correlation structure.

| Analysis Type | Test | Nominal Level | Logistic Mixed Model |  |  | Liability Threshold Model |  |  |
| --- | --- | --- | --- | --- | --- | --- | --- | --- |
|  |  |  | Random | Baseline | Sum | Random | Baseline | Sum |
| Prospective | GEE(ind) | 0.05 | **$5.38 \times 10^{-2}$** | **$5.08 \times 10^{-2}$** | **$5.27 \times 10^{-2}$** | **$5.36 \times 10^{-2}$** | **$5.19 \times 10^{-2}$** | **$5.38 \times 10^{-2}$** |
|  |  | 0.01 | **$1.18 \times 10^{-2}$** | **$1.04 \times 10^{-2}$** | **$1.13 \times 10^{-2}$** | **$1.17 \times 10^{-2}$** | **$1.07 \times 10^{-2}$** | **$1.17 \times 10^{-2}$** |
|  |  | 0.001 | **$1.32 \times 10^{-3}$** | **$1.16 \times 10^{-3}$** | **$1.23 \times 10^{-3}$** | **$1.37 \times 10^{-3}$** | **$1.14 \times 10^{-3}$** | **$1.37 \times 10^{-3}$** |
|  |  | 0.0001 | **$1.67 \times 10^{-4}$** | **$1.28 \times 10^{-4}$** | **$1.43 \times 10^{-4}$** | **$1.34 \times 10^{-4}$** | **$1.36 \times 10^{-4}$** | **$1.76 \times 10^{-4}$** |
| | GEE(AR1) | 0.05 | **$5.36 \times 10^{-2}$** | $5.02 \times 10^{-2}$ | **$5.26 \times 10^{-2}$** | **$5.34 \times 10^{-2}$** | **$5.17 \times 10^{-2}$** | **$5.37 \times 10^{-2}$** |
|  |  | 0.01 | **$1.16 \times 10^{-2}$** | **$1.04 \times 10^{-2}$** | **$1.12 \times 10^{-2}$** | **$1.16 \times 10^{-2}$** | **$1.06 \times 10^{-2}$** | **$1.17 \times 10^{-2}$** |
|  |  | 0.001 | **$1.31 \times 10^{-3}$** | **$1.13 \times 10^{-3}$** | **$1.21 \times 10^{-3}$** | **$1.36 \times 10^{-3}$** | **$1.14 \times 10^{-3}$** | **$1.36 \times 10^{-3}$** |
| | | 0.0001 | **$1.73 \times 10^{-4}$** | $1.19 \times 10^{-4}$ | **$1.37 \times 10^{-4}$** | **$1.32 \times 10^{-4}$** | **$1.35 \times 10^{-4}$** | **$1.78 \times 10^{-4}$** |
|  | GEE(mix) | 0.05 | **$5.34 \times 10^{-2}$** | **$5.07 \times 10^{-2}$** | **$5.26 \times 10^{-2}$** | **$5.34 \times 10^{-2}$** | **$5.19 \times 10^{-2}$** | **$5.37 \times 10^{-2}$** |
|  |  | 0.01 | **$1.17 \times 10^{-2}$** | **$1.04 \times 10^{-2}$** | **$1.13 \times 10^{-2}$** | **$1.16 \times 10^{-2}$** | **$1.07 \times 10^{-2}$** | **$1.17 \times 10^{-2}$** |
|  |  | 0.001 | **$1.29 \times 10^{-3}$** | **$1.17 \times 10^{-3}$** | **$1.22 \times 10^{-3}$** | **$1.38 \times 10^{-3}$** | **$1.14 \times 10^{-3}$** | **$1.36 \times 10^{-3}$** |
|  |  | 0.0001 | **$1.70 \times 10^{-4}$** | **$1.29 \times 10^{-4}$** | **$1.37 \times 10^{-4}$** | **$1.31 \times 10^{-4}$** | **$1.30 \times 10^{-4}$** | **$1.70 \times 10^{-4}$** |
| | GMMAT | 0.05 | $3.89 \times 10^{-2}$ | $3.53 \times 10^{-2}$ | $4.76 \times 10^{-2}$ | $4.80 \times 10^{-2}$ | $4.89 \times 10^{-2}$ | $4.91 \times 10^{-2}$ |
| | | 0.01 | $6.07 \times 10^{-3}$ | $5.24 \times 10^{-3}$ | $9.08 \times 10^{-3}$ | $9.29 \times 10^{-3}$ | $9.51 \times 10^{-3}$ | $9.33 \times 10^{-3}$ |
| | | 0.001 | $4.29 \times 10^{-4}$ | $3.74 \times 10^{-4}$ | $7.84 \times 10^{-4}$ | $8.63 \times 10^{-4}$ | $8.96 \times 10^{-4}$ | $8.33 \times 10^{-4}$ |
| | | 0.0001 | $2.20 \times 10^{-5}$ | $2.20 \times 10^{-5}$ | $6.80 \times 10^{-5}$ | $6.30 \times 10^{-5}$ | $9.10 \times 10^{-5}$ | $8.80 \times 10^{-5}$ |
| Retrospective | L-BRAT(ind) | 0.05 | $4.93 \times 10^{-2}$ | $4.91 \times 10^{-2}$ | $4.98 \times 10^{-2}$ | $5.01 \times 10^{-2}$ | $4.99 \times 10^{-2}$ | $4.98 \times 10^{-2}$ |
| | | 0.01 | $9.45 \times 10^{-3}$ | $9.60 \times 10^{-3}$ | $9.84 \times 10^{-3}$ | $9.90 \times 10^{-3}$ | $9.75 \times 10^{-3}$ | $9.55 \times 10^{-3}$ |
| | | 0.001 | $8.30 \times 10^{-4}$ | $9.78 \times 10^{-4}$ | $9.24 \times 10^{-4}$ | $9.55 \times 10^{-4}$ | $9.45 \times 10^{-4}$ | $8.78 \times 10^{-4}$ |
| | | 0.0001 | $7.20 \times 10^{-5}$ | $9.50 \times 10^{-5}$ | $8.20 \times 10^{-5}$ | $8.20 \times 10^{-5}$ | $9.40 \times 10^{-5}$ | $9.20 \times 10^{-5}$ |
| | L-BRAT(AR1) | 0.05 | $4.93 \times 10^{-2}$ | $4.88 \times 10^{-2}$ | $4.97 \times 10^{-2}$ | $4.99 \times 10^{-2}$ | $4.98 \times 10^{-2}$ | $4.97 \times 10^{-2}$ |
| | | 0.01 | $9.48 \times 10^{-3}$ | $9.72 \times 10^{-3}$ | $9.78 \times 10^{-3}$ | $9.84 \times 10^{-3}$ | $9.76 \times 10^{-3}$ | $9.55 \times 10^{-3}$ |
| | | 0.001 | $8.26 \times 10^{-4}$ | $9.62 \times 10^{-4}$ | $9.22 \times 10^{-4}$ | $9.17 \times 10^{-4}$ | $9.47 \times 10^{-4}$ | $8.48 \times 10^{-4}$ |
| | | 0.0001 | $8.80 \times 10^{-5}$ | $9.60 \times 10^{-5}$ | $8.20 \times 10^{-5}$ | $7.10 \times 10^{-5}$ | $1.02 \times 10^{-4}$ | $8.90 \times 10^{-5}$ |
| | L-BRAT(mix) | 0.05 | $4.93 \times 10^{-2}$ | $4.91 \times 10^{-2}$ | $4.99 \times 10^{-2}$ | $5.01 \times 10^{-2}$ | $4.98 \times 10^{-2}$ | $4.98 \times 10^{-2}$ |
| | | 0.01 | $9.57 \times 10^{-3}$ | $9.61 \times 10^{-3}$ | $9.86 \times 10^{-3}$ | $9.88 \times 10^{-3}$ | $9.79 \times 10^{-3}$ | $9.54 \times 10^{-3}$ |
| | | 0.001 | $8.35 \times 10^{-4}$ | $9.86 \times 10^{-4}$ | $9.26 \times 10^{-4}$ | $9.57 \times 10^{-4}$ | $9.37 \times 10^{-4}$ | $8.78 \times 10^{-4}$ |
| | | 0.0001 | $8.20 \times 10^{-5}$ | $1.01 \times 10^{-4}$ | $8.60 \times 10^{-5}$ | $7.40 \times 10^{-5}$ | $9.70 \times 10^{-5}$ | $8.90 \times 10^{-5}$ |
| | RGMMAT | 0.05 | $4.72 \times 10^{-2}$ | $4.91 \times 10^{-2}$ | $4.98 \times 10^{-2}$ | $4.93 \times 10^{-2}$ | $4.99 \times 10^{-2}$ | $4.98 \times 10^{-2}$ |
| | | 0.01 | $8.76 \times 10^{-3}$ | $9.64 \times 10^{-3}$ | $9.85 \times 10^{-3}$ | $9.63 \times 10^{-3}$ | $9.78 \times 10^{-3}$ | $9.55 \times 10^{-3}$ |
| | | 0.001 | $7.20 \times 10^{-4}$ | $9.52 \times 10^{-4}$ | $9.09 \times 10^{-4}$ | $9.12 \times 10^{-4}$ | $9.43 \times 10^{-4}$ | $8.75 \times 10^{-4}$ |
| | | 0.0001 | $6.80 \times 10^{-5}$ | $8.90 \times 10^{-5}$ | $8.20 \times 10^{-5}$ | $7.70 \times 10^{-5}$ | $9.10 \times 10^{-5}$ | $9.30 \times 10^{-5}$ |

To compare power, we considered five effect sizes at the two causal SNPs, and tested association between the trait and the first causal SNP. Empirical power was calculated at the significance level 10^−3^, based on 1,000 simulated replicates. Figure 1 demonstrates the power results for each method. In all the simulation settings, the retrospective tests consistently had higher power than the prospective tests. The L-BRAT association tests under three different working correlation structures had similar power. The RGMMAT method also achieved high power. In contrast, the prospective GEE methods had the lowest power in all settings except under the baseline sampling and the liability threshold model, in which GMMAT performed the worst in power. Overall, we found that the baseline sampling scheme generated the highest power under different trait models, while the sum sampling scheme had a power gain over the random sampling scheme under the logistic mixed model, but was less powerful under the liability threshold model. These results suggest that L-BRAT and RGMMAT outperform the prospective tests, and the power of L-BRAT is not sensitive to the choice of the working correlation structure.

**Figure 1.**
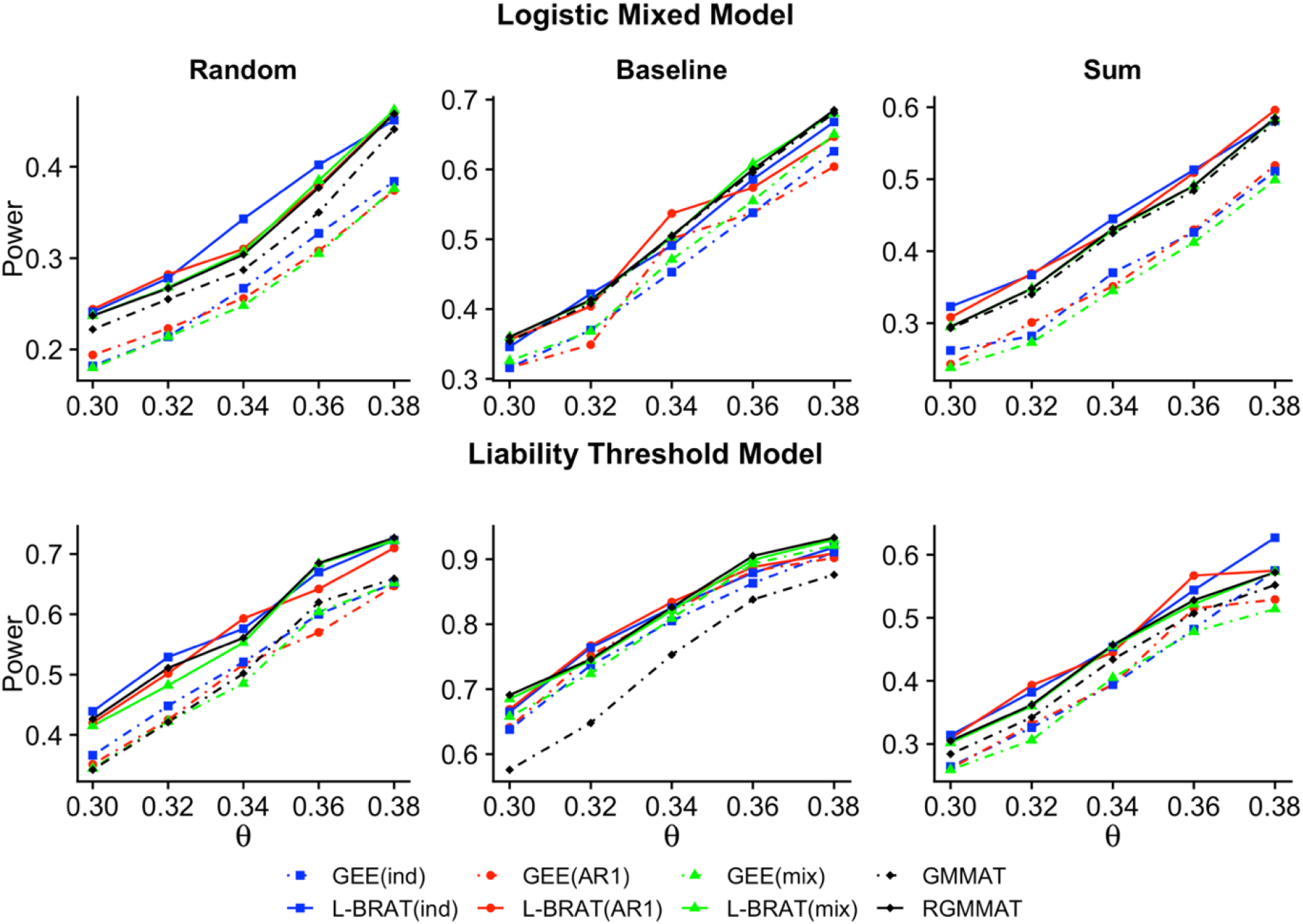
Empirical power of L-BRAT, RGMMAT, GEE, and GMMAT. Power is based on 1,000 simulated replicates at five time points with *α* = 10 ^−3^. In the upper panel, the trait is simulated by the logistic mixed model, and in the lower panel, it is by the liability threshold model. Power results are demonstrated in samples of 2,000 individuals according to three different ascertainment schemes: random, baseline, and sum. This figure appears in color in the electronic version of this article.

## 4. Analysis of Longitudinal Cocaine Use Data from VACS

We illustrated the utility of our proposed methods by analyzing a GWAS dataset of cocaine use from VACS (Justice et al. 2006). VACS is a multi-center, longitudinal observational study of HIV infected and uninfected veterans whose primary objective is to understand the risk of alcohol and other substance abuse in individuals with HIV infection. We analyzed longitudinal cocaine use in patient surveys collected at six clinic visits on 2,470 participants. Among them, 69.8% are African Americans (AAs), 19.3% are European Americans (EAs), and 10.9% are of other races. We considered the responses at each visit as 0 if individuals had never tried cocaine or had not used cocaine in the last year, and as 1 if individuals had used cocaine in the last year. The proportion of case subjects at each visit ranges from 13.7% (*n* = 192) to 24.3% (*n* = 526), and the missing rate at each visit ranges from 3.0% to 44.2%.

All samples were genotyped on the Illumina OmniExpress BeadChip. After data cleaning, there are 2,458 individuals available for genotype imputation. IMPUTE2 (Howie et al. 2009) was used for imputation using the 1000 Genomes Phase 3 data as a reference panel. We excluded subjects who did not meet either of the following criteria: (1) completeness (i.e., proportion of successfully imputed SNPs) > 95% and (2) empirical self-kinship < 0.525 (i.e., empirical inbreeding coefficient < 0.05). Based on the above criteria, 2,231 individuals were retained in the analysis, with 2,114 males and 117 females, of whom 1,557 are AAs, 431 are EAs, and 243 are of other races. There are 1,433 individuals who had never used cocaine during the study period, 639 individuals who sometimes used cocaine, i.e., exhibited response variation, and 159 individuals who had used cocaine at least once every year over the course of the study. SNPs that satisfied all of the following quality-control conditions were included in the analysis: (1) call rate > 95%, (2) Hardy-Weinberg *χ*^2^ statistic P-value > 10^−6^, and (3) MAF > 1%. All together there are a final set of 10,215,072 SNPs retained in the analysis.

### 4.1. Association Analysis

Genome-wide association testing for longitudinal cocaine use was performed using L-BRAT, RGMMAT, and the prospective GEE and GMMAT tests in the entire VACS sample. Sex, age at baseline, HIV status, and time were included as covariates in the analysis. The top ten principal components (PCs) that explained 89.4% of the total genetic variation were included as covariates to control for population structure. We considered two working correlation structures: independent and AR(1). For the L-BRAT and RGMMAT methods, the GRM was calculated using the LD pruned SNPs with MAF > 0.05.

To compare the performance of longitudinal association analysis with that of univariate analysis on the summary metrics of cocaine use in VACS, we considered two alternative cocaine phenotypes: baseline and trajectories. Longitudinal cocaine use trajectories were obtained using a growth mixture model that clusters longitudinal data into discrete growth trajectory curves (Muthén 2004). We fit a logistic model with a polynomial function of time. The number of groups was chosen based on the Bayesian information criterion (BIC). Each individual was then assigned to the trajectory with the highest probability of membership. Figure 2 shows the four cocaine use trajectory groups identified in the VACS sample. They were labeled as mostly never (0, *n* = 1,682), moderate decrease (1, *n* = 296), elevated chronic (2, *n* = 86), and mostly frequent (3, *n* = 167). We used CARAT, a case-control retrospective association test (Jiang et al. 2016), for the analysis of cocaine use at baseline, adjusted for sex, age at baseline, and HIV status. Cumulative logit model was used to test for association between the four ordered cocaine use trajectory groups and each of the SNPs, with adjustment for sex, age at baseline, HIV status, and the top ten PCs.

**Figure 2.**
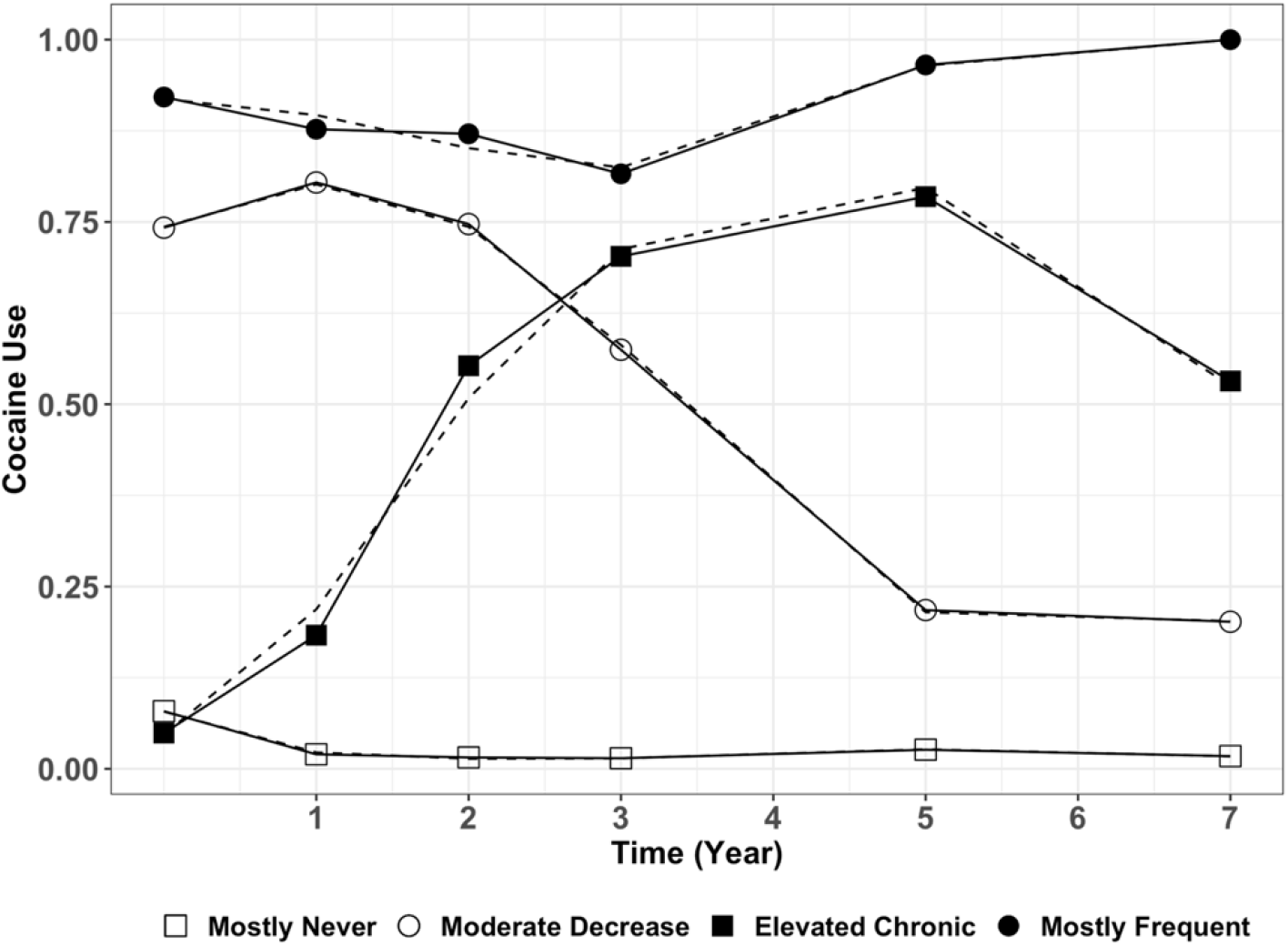
Group-based cocaine use trajectories in VACS. Dashed lines represent the estimated trajectories, solid lines represent the observed mean cocaine use for each trajectory group. Time is the number of years since the baseline visit.

None of the retrospective tests exhibited evidence of inflation in the quantile-quantile (Q-Q) plot (Web Figure 1). The genomic control inflation factors were 0.993 and 0.991 for the L-BRAT genome scan under the independent and AR(1) working correlation, respectively, and 0.984 for the RGMMAT analysis. The prospective GEE tests showed some evidence of deflation in the Q-Q plot. The genomic control factors were 0.938 and 0.937 for the GEE tests under the independent and AR(1) working correlation. The most conservative test was GMMAT, with a genomic control factor 0.802.

Table 2 reports the results for SNPs for which at least one of the longitudinal tests gives a P-value < 2 × 10^−7^. Among them, the L-BRAT tests produced the smallest P-values, RGMMAT and the trajectory-based analysis had comparable results, while GEE, GMMAT, and CARAT generated much larger P-values. The Manhattan plot of the smallest P-value from these tests in the VACS cocaine use data is shown in Web Figure 2. Among the top SNPs listed in Table 2, there are two SNPs, rs551879660 and rs150191017, located at 3p12 and 13q12 respectively, that reach the genome-wide significance (P = 2.00 × 10^−8^ and 3.77 × 10^−8^, respectively). Each of these SNPs was reported to have MAF < 1% in the 1000 Genomes (MAF = 0.68% and 0.98%, respectively). The MAFs of the two SNPs were 1.2% and 1.1% in the entire VACS sample, respectively, and were slightly higher in the AA sample (MAF = 1.6% and 1.5%, respectively). Although both SNPs have MAF > 1%, given the small sample size of VACS, there is limited information on them. SNP rs150191017 is located 31.5 kb from the gene *AL161616.2* which was reported to be associated with venlafaxine treatment response in a generalized anxiety disorder GWAS (Jung et al. 2017). A cluster of five SNPs in LD, rs76386683, rs114386843, rs186274502, rs376616438, and rs187855416, located at 9q33, showed association with longitudinal cocaine use (P = 1.85 × 10^−7^ − 1.93 × 10^−7^). They are near *OR1L4*, an olfactory receptor gene that was reported to be associated with major depressive disorder (Wong et al. 2017). A cluster of olfactory receptor genes between *OR3A1* and *OR3A2* that belong to the olfactory receptor gene family were identified in a recent GWAS of cocaine dependence and related traits (Gelernter et al. 2014). The other three SNPs, rs188222191, rs1014278, rs75132056, are located at 5q21 (P = 1.28 × 10^−7^, 1.43 × 10^−7^ and 8.92 × 10^−8^, respectively), close to the gene *EFNA5*, which was identified in several GWAS to be associated with bipolar disorder and schizophrenia (Wang et al. 2010). There was also evidence of association with rs114629793 (P = 8.65 × 10^−8^). This SNP is in an intron of the gene encoding *PSD3*, located at 8p22. Recently, two schizophrenia GWAS have identified association between *PSD3* and schizophrenia (Goes et al. 2015; Li et al. 2017b), and one study has shown that *PSD3* is associated with paliperidone treatment response in schizophrenic patients (Li et al. 2017a). Gene network analysis revealed that *PSD3* is one of the differentially expressed hub genes that involve dysfunction of brain reward circuitry in cocaine use disorder (Ribeiro et al. 2017).

**Table 2.** SNPs with P-value < 2×10^−7^ in at least one of the longitudinal tests in the entire VACS sample. The smallest P-value among all tests at the given SNPs are in bold. ^a^ CARAT applied to cocaine use at baseline, ^b^ Cumulative logit model applied to the four ordered cocaine use trajectory group.

| Chr | Gene Region | SNP | Position | MAF | GEE (ind) | GEE (AR1) | GMMAT | L-BRAT (ind) | L-BRAT (AR1) | RGMMAT | CARAT<sup>a</sup> (BL) | CL<sup>b</sup> (traj) |
| --- | --- | --- | --- | --- | --- | --- | --- | --- | --- | --- | --- | --- |
| 3 | *NIPA2P2* | rs551879660 | 75,146,492 | 0.012 | $1.87 \times 10^{-4}$ | $7.14 \times 10^{-4}$ | $9.07 \times 10^{-4}$ | **$2.00 \times 10^{-8}$** | $3.19 \times 10^{-6}$ | $4.13 \times 10^{-5}$ | $5.78 \times 10^{-4}$ | $3.35 \times 10^{-5}$ |
| 5 | *EFNA5* | rs188222191 | 105,411,547 | 0.042 | $6.86 \times 10^{-6}$ | $1.65 \times 10^{-5}$ | $8.87 \times 10^{-5}$ | **$1.28 \times 10^{-7}$** | $4.17 \times 10^{-7}$ | $2.69 \times 10^{-6}$ | $8.95 \times 10^{-5}$ | $2.72 \times 10^{-5}$ |
| | | rs1014278 | 105,471,506 | 0.057 | $1.02 \times 10^{-5}$ | $1.10 \times 10^{-5}$ | $1.24 \times 10^{-4}$ | $1.50 \times 10^{-7}$ | **$1.43 \times 10^{-7}$** | $4.88 \times 10^{-6}$ | $5.94 \times 10^{-5}$ | $3.00 \times 10^{-5}$ |
| | | rs75132056 | 105,480,442 | 0.05 | $1.05 \times 10^{-5}$ | $2.42 \times 10^{-5}$ | $1.89 \times 10^{-4}$ | **$8.92 \times 10^{-8}$** | $2.89 \times 10^{-7}$ | $8.55 \times 10^{-6}$ | $2.59 \times 10^{-4}$ | $2.31 \times 10^{-5}$ |
| 8 | *PSD3* | rs114629793 | 18,403,754 | 0.012 | $3.12 \times 10^{-4}$ | $4.73 \times 10^{-4}$ | $1.44 \times 10^{-4}$ | **$8.65 \times 10^{-8}$** | $3.60 \times 10^{-7}$ | $2.82 \times 10^{-6}$ | $5.12 \times 10^{-4}$ | $3.06 \times 10^{-6}$ |
| 9 | *OR1L4* | rs76386683 | 125,467,023 | 0.012 | $1.48 \times 10^{-4}$ | $9.15 \times 10^{-5}$ | $2.86 \times 10^{-4}$ | $1.03 \times 10^{-6}$ | **$1.93 \times 10^{-7}$** | $5.92 \times 10^{-6}$ | $4.80 \times 10^{-4}$ | $3.30 \times 10^{-6}$ |
| | | rs114386843 | 125,469,425 | 0.012 | $1.47 \times 10^{-4}$ | $9.05 \times 10^{-5}$ | $2.82 \times 10^{-4}$ | $1.01 \times 10^{-6}$ | **$1.88 \times 10^{-7}$** | $5.78 \times 10^{-6}$ | $4.75 \times 10^{-4}$ | $3.22 \times 10^{-6}$ |
| | | rs186274502 | 125,471,416 | 0.012 | $1.47 \times 10^{-4}$ | $9.05 \times 10^{-5}$ | $2.82 \times 10^{-4}$ | $1.01 \times 10^{-6}$ | **$1.88 \times 10^{-7}$** | $5.78 \times 10^{-6}$ | $4.75 \times 10^{-4}$ | $3.22 \times 10^{-6}$ |
| | | rs376616438 | 125,472,267 | 0.012 | $1.44 \times 10^{-4}$ | $8.95 \times 10^{-5}$ | $2.77 \times 10^{-4}$ | $9.79 \times 10^{-7}$ | **$1.85 \times 10^{-7}$** | $5.62 \times 10^{-6}$ | $4.79 \times 10^{-4}$ | $3.20 \times 10^{-6}$ |
| | | rs187855416 | 125,474,459 | 0.012 | $1.44 \times 10^{-4}$ | $8.95 \times 10^{-5}$ | $2.77 \times 10^{-4}$ | $9.79 \times 10^{-7}$ | **$1.85 \times 10^{-7}$** | $5.62 \times 10^{-6}$ | $4.79 \times 10^{-4}$ | $3.20 \times 10^{-6}$ |
| 11 | *AP000851.1* | rs139780693 | 102,509,700 | 0.03 | $2.60 \times 10^{-5}$ | $1.04 \times 10^{-5}$ | $2.78 \times 10^{-4}$ | $5.83 \times 10^{-7}$ | **$1.26 \times 10^{-7}$** | $1.35 \times 10^{-5}$ | $1.06 \times 10^{-4}$ | $2.00 \times 10^{-6}$ |
| 13 | *AL161616.2* | rs150191017 | 31,962,649 | 0.011 | $4.26 \times 10^{-5}$ | $9.72 \times 10^{-5}$ | $7.32 \times 10^{-5}$ | **$3.77 \times 10^{-8}$** | $3.09 \times 10^{-7}$ | $7.87 \times 10^{-7}$ | $3.74 \times 10^{-4}$ | $5.48 \times 10^{-7}$ |

We further analyzed the data separately in each population, adjusted for the top ten PCs obtained within the group, and then combined the results from the three groups by meta-analysis using the optimal weights for score statistics that have essentially the same power as the inverse variance weighting (Zhou et al. 2011). Web Table 1 gives the results in the 1,557 AA samples. All the top twelve SNPs listed in Table 2 had a P-value < 5 × 10^−5^ in at least one of the longitudinal tests in AAs. L-BRAT consistently gave the smallest P-values among all the longitudinal tests. The results from the three groups (AAs, EAs and other races) were combined by meta-analysis. The meta-analysis P-values were of the same order of magnitude as that obtained from the entire sample adjusted for population structure for each longitudinal test (Table 3). All the top twelve SNPs listed in Table 2 had a meta-analysis P-value < 8 × 10^−7^ in at least one of the longitudinal tests. Among them, the L-BRAT test with either an independent or AR(1) working correlation gave the smallest meta-analysis P-values.

**Table 3.**
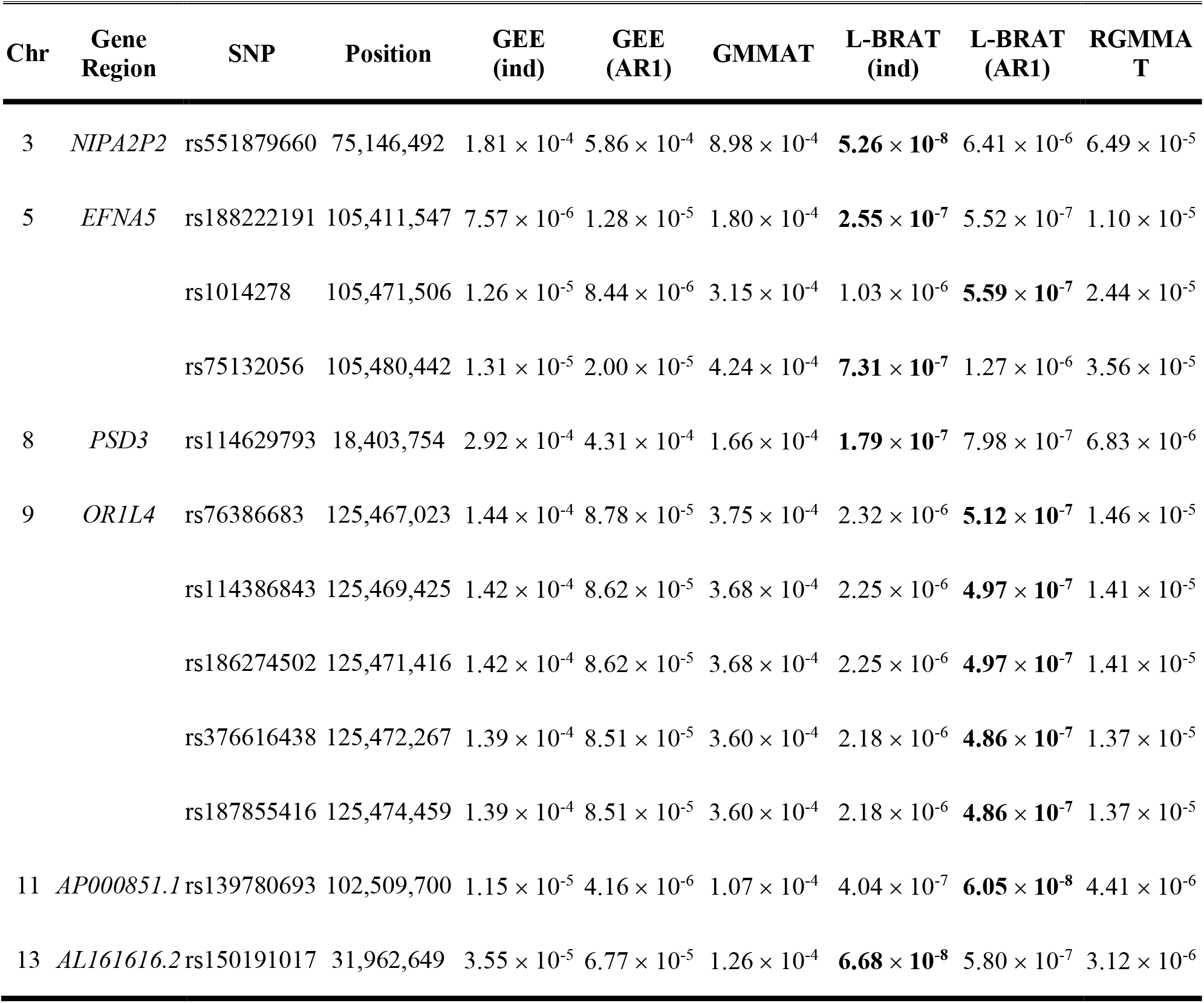
Meta-analysis results of the top twelve SNPs from Table 2 in the VACS data. The smallest P-value among all tests at the given SNPs are in bold.

### 4.2. Pathway Analysis

We then performed pathway analysis on the top SNPs for which at least one of the longitudinal tests had a P-value < 5 × 10^−5^ using the Ingenuity Pathway Analysis (IPA). We identified two significant canonical pathways that belong to the neurotransmitters and nervous system signaling. The first one is the opioid signaling pathway (P = 1.41 × 10^−4^, adjusted P = 0.010), which plays an important role in opioid tolerance and dependence. Studies have shown that chronic administration of cocaine and opioids are associated with changes in dopamine transporters and opioid receptors in various brain regions (Le Merrer et al. 2009; Soderman and Unterwald 2009). The second significant pathway is the axonal guidance signaling pathway (P = 2.54 × 10^−4^, adjusted P = 0.012), which is critical for neural development. The mRNA expression levels of axon guidance molecules have been found to be altered in some brain regions of cocaine-treated rats, which may contribute to drug abuse-associated cognitive impairment (Bahi and Dreyer 2005; Jassen 2006). Each of the two pathways remained significant when we performed pathway analysis, using the same P-value cutoff value to select top SNPs, based on the L-BRAT results generated under the independence and AR(1) working correlation, respectively. In contrast, only the opioid signaling pathway was significant based on the results from the GEE analysis using the independent working correlation, and only the axonal guidance signaling pathway was significant based on the RGMMAT results, whereas neither of them remained significant based on the GMMAT results and that from the GEE analysis with an AR(1) working correlation. These results demonstrate that L-BRAT provides more informative association results to help identify biological relevant pathways.

### 4.3. eQTL Enrichment Analysis

Lastly, we performed an enrichment analysis to see whether the top SNPs in our analysis are more likely to regulate brain gene expression. We considered the local expression quantitative trait loci (*cis*-eQTLs) reported in 13 human brain regions from the Genotype-Tissue Expression (GTEx) project (GTEx Consortium 2013, 2017), including amygdala, anterior cingulate cortex, caudate, cerebellar hemisphere, cerebellum, cortex, frontal cortex, hippocampus, hypothalamus, nucleus accumbens, putamen, spinal cord, and substantia nigra. Fisher’s exact test was used to assess the enrichment of eQTLs (FDR < 0.05) in the top 2,778 SNPs for which at least one of the longitudinal tests had a P-value < 10^−4^ in the VACS sample. Among the 13 brain regions, amygdala is the only region in which eQTLs showed significant enrichment in our top SNP list (odds ratio = 2.06, P = 3.0 × 10^−5^).

### 4.4. Replication of Top Findings

We used an independent cocaine dependence case-control GWAS from the Yale-Penn study (Gelernter et al. 2014) to replicate the top findings from our longitudinal analysis results in VACS. The summary statistics from the Yale-Penn cocaine dependence GWAS were obtained. Note that the lifetime cocaine dependence diagnosis was made using the Semi-Structured Assessment for Drug Dependence and Alcoholism (SSADDA) (Pierucci-Lagha et al. 2005) which is different from the outcome used in VACS, and there were no longitudinal phenotype measures in Yale-Penn. Nevertheless, we performed pathway analysis using the SNP summary statistics of Yale-Penn to replicate the two pathways identified in the VACS sample. Among the top 2,778 SNPs for which at least one of the longitudinal tests had a P-value < 10^−4^, we were able to retrieve 2,602 SNP summary statistics from Yale-Penn. Pathway analysis was conducted on the top 84 SNPs that had a P-value < 0.05. Although none of the top twelve SNPs in Table 2 had a P-value < 0.05 in the Yale-Penn AA sample, each of the two pathways remained significant: the opioid signaling pathways (P = 5.67 × 10^−4^, adjusted P = 3.77 × 10^−3^) and the axonal guidance signaling (P = 2.89 × 10^−4^, adjusted P = 2.97 × 10^−3^).

## 5. Discussion

Longitudinal data can be used in GWAS to improve power for identification of genetic variants and environmental factors that influence complex traits over time. In this study, we have developed L-BRAT, a retrospective association testing method for longitudinal binary outcomes. L-BRAT is based on GEE, thus it requires assumptions on the mean but not the full distribution of the outcome. Correct specification of the covariance of repeated measurements within each individual is not required, instead, a working covariance matrix is assumed. The significance of the L-BRAT association test is assessed retrospectively by considering the conditional distribution of the genotype at the variant of interest, given phenotype and covariate information, under the null hypothesis of no association. Features of L-BRAT include the following: (1) it is computationally feasible for genetic studies with millions of variants, (2) it allows both static and time-varying covariates to be included in the analysis, (3) it allows different individuals to have measurements at different time points, and (4) it has correct type I error in the presence of ascertainment and trait model misspecification. For comparison, we also propose a retrospective, logistic mixed model-based association test, RGMMAT, which requires specification of the full distribution of the outcome. Random effects are used to model dependence among observations for an individual. Like L-BRAT, RGMMAT is a retrospective analysis in which genotypes are treated as random conditional on the phenotype and covariates. As a result, RGMMAT is also robust to misspecification of the model for the phenotype distribution.

Through simulation, we demonstrated that the type I error of L-BRAT was well calibrated under different trait models and ascertainment schemes, whereas the type I error of the prospective GEE method was inflated relative to nominal levels. In the GLMM-based methods, GMMAT, a prospective analysis, was overly conservative, whereas the retrospective version, RGMMAT, was able to maintain correct type I error. We further demonstrated that the two retrospective tests, L-BRAT and RGMMAT, provided higher power to detect association than the prospective GEE and GMMAT tests under all the trait models and ascertainment schemes considered in the simulations. The choice of the working correlation matrix in L-BRAT resulted in little loss of power. We applied L-BRAT and RGMMAT to longitudinal association analysis of cocaine use in the VACS data, where we identified six novel genes that are associated with cocaine use. Moreover, our pathway analysis identified two significant pathways associated with longitudinal cocaine use: the opioid signaling pathway and the axonal guidance signaling pathway. We were able to replicate both pathways in a cocaine dependence case-control GWAS from the Yale-Penn study. Lastly, we illustrated that the top SNPs identified by our methods are more likely to be the amygdala eQTLs in the GTEx data. The amygdala plays an important role in the processing of memory, decision-making, and emotional responses, and contributes to drug craving that leads to addiction and relapse (Hyman and Malenka 2001; Warlow et al. 2017). These findings verify that L-BRAT is able to detect important loci in a genome scan and to provide novel insights into the disease mechanism in relevant tissues.

The L-BRAT and RGMMAT methods are designed for single-variant association analysis of longitudinally measured binary outcomes. However, single-variant association tests in general have limited power to detect association for low-frequency or rare variants in sequencing studies. We have previously developed longitudinal burden test and sequence kernel association test, LBT and LSKAT, to analyze rare variants with longitudinal quantitative phenotypes (Wang et al. 2017). Both tests are based on a prospective approach. To extend L-BRAT and RGMMAT to rare variant analysis with longitudinal binary data, we could consider either a linear statistic or a quadratic statistic that combines the retrospective score test at each variant in a gene region. In addition, the genetic effect in L-BRAT and RGMMAT is assumed to be constant. We could consider an extension to allow for time-varying genetic effect so that the fluctuation of genetic contributions to the trait value over time is well calibrated.

## ACKNOWLEDGEMENTS

This study was supported by the National Institute of Health grants K01 AA023321 and R21 AA022870. The authors appreciate the support of the Veterans Aging Cohort Study and Yale Center for Genome Analysis.

## SUPPORTING INFORMATION

Additional supporting information referenced in Section 4 may be found online in the Supporting Information section at the end of the article. R package implementing L-BRAT and RGMMAT can be found at https://github.com/ZWang-Lab/LBRAT.

